# Using QSARs for predictions in drug delivery

**DOI:** 10.1101/727172

**Authors:** Edgardo Rivera-Delgado, Alison Xin, Horst A. von Recum

**Affiliations:** Department of Biomedical Engineering, Case Western Reserve University, Cleveland, Ohio, United States of America; Science Research and Engineering Program, Hathaway Brown School, Shaker Heights, Ohio

## Abstract

Drug delivery research is an inherently empirical process, however high-throughput approaches could take advantage of understanding drug/material interactions such as from electrostatic, hydrophobic, or other non-covalent interactions between therapeutic molecules and a drug delivery polymer. Cyclodextrin polymers have been investigated for drug delivery specifically due to their capacity to exploit this affinity interaction to change the rate of drug release. Testing drug candidates; however, for affinity is time-consuming, making computational predictions more effective. One option, molecular "docking" programs, provide predictions of affinity, but lack reliability, as their accuracy with cyclodextrin remains unverified experimentally. Alternatively, quantitative structure-activity relationship models (QSARs), which analyze statistical relationships between molecular properties, appear more promising. Previously constructed QSARs for cyclodextrin are not publicly available, necessitating an openly accessible model. Around 600 experimental affinities between cyclodextrin and guest molecules were cleaned and imported from published research. The software PaDEL-Descriptor calculated over 1000 chemical descriptors for each molecule, which were then analyzed in R to create several QSARs with different statistical methods. These QSARs proved highly time efficient, calculating in minutes what docking programs would take hours to accomplish. Additionally, on test sets, QSARs reached R^2^ values of around 0.7-0.8. The speed, accuracy, and accessibility of these QSARs improve evaluation of individual drugs and facilitate screening of large datasets for potential candidates in cyclodextrin affinity-based delivery systems. An app was built to rapidly access model predictions for end users using the "shiny" library in R. To demonstrate the usability for drug release planning, the QSAR predictions were coupled with a mechanistic model of diffusion within the app. Integrating new modules should provide an accessible approach to use other cheminformatic tools in the field of drug delivery.

## 1. Introduction

Drug delivery is an inherently challenging process to predict, due to situational constraints (device size, polymer properties, diffusivity). Our lab and others have explored use of polymers containing specific chemistries allowing stronger polymer drug interactions to control the rate of drug release. As these affinities are increased the dependence on other factors such as size and diffusivity decrease. This field is termed "Affinity-based drug delivery", and relies on interactions between a drug delivery system and drug molecules, improves effectiveness of medication by extending the duration of drug release and thereby lengthening the duration of the treatment[1]. Mathematical modeling of these affinity systems has shown that the strength of the affinity interaction, the ratio of host binding sites to guest ligands, and the molecular path length of diffusion influence the transport of molecules out of the system. Of these physical forces, the affinity strength plays an important role in the classification of the system and the timescale of drug release[2]. Affinity interaction can be associated with a variety of physical properties, including charge, hydrophobicity, Van der Waals forces, etc. In the fields of biomaterials and drug delivery, affinity delivery has been used with small molecule drugs[3], proteins[1], cytokines, and antibodies[4].

Our lab tests rings of glucose molecules as affinity hosts called cyclodextrins, which are particularly promising affinity drug delivery hosts due to their structural properties, biocompatibility and versatility. The most common cyclodextrin are composed of a ring 6, 7, or 8 glucose molecules (α, β, and γ-cyclodextrin, respectively), and the conformation of the hydroxyl groups of the ring create a basket-like structure with a hydrophobic interior and hydrophilic interior, allowing for complexation with drug molecules (Figure 1). Additionally, cyclodextrin can be polymerized into a variety of materials, including microparticles, viscous gels, and solid films. Unfortunately, experiments to confirm sustained release from the affinity guest-host system often takes weeks, making testing large numbers of potential candidates for cyclodextrin release systems impractical.

**Fig 1.**
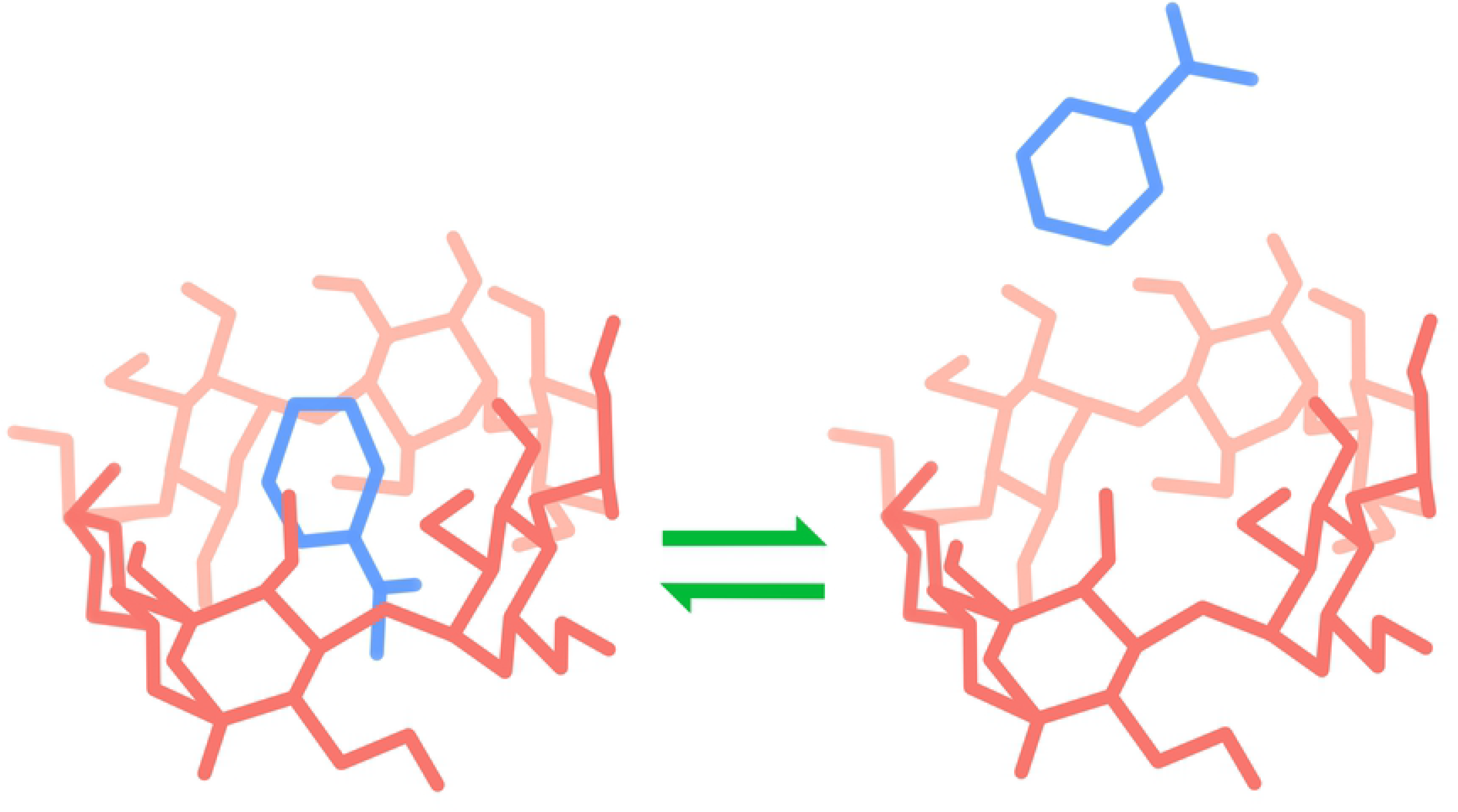
Cyclodextrin complexation.

As an alternative to experimental testing, candidate molecules can be analyzed computationally. Predicting the binding affinity between cyclodextrin and drug molecules allows for the processing of molecules on the scale of minutes rather than weeks. There are two major methods for predicting molecular interaction: docking models and QSARs. Docking models use molecular force fields, which simulate interactions and potential energy between atoms. Force field parameters may be derived from experiments, calculations from quantum mechanics, or both[5]. In addition to providing a numeric estimate for binding affinity, docking programs produce visualizations of how molecules interact. QSARs, or Quantitative Structure-Activity Relationship models, statistically predict molecular interactions using molecular descriptors. Molecular descriptors are certain physical or chemical characteristics of molecules that can be evaluated numerically (for example, the number of hydrogen atoms or the length of the longest bond chain). Many different types of regression models and statistical learning methods can be used as QSARs, ranging in complexity from linear models to artificial neural networks[6].

Previous investigations have been made on the accuracy of both docking and QSARs in predicting cyclodextrin affinity, but examining a sample of these papers reveals several concerns (Table 1). Notably, all of the investigated models used software hidden behind a paywall or only available with a license[7–14]. Additionally, many models lacked proper verification. Following Tropsha’s publication detailing best practices for QSAR development, a completely verified model should undergo leave-one-out cross-validation (LOO-CV) (reported as Q^2^), y-randomization, pass a variety of internal accuracy tests, and be analyzed for applicability domain. Additionally, models should be evaluated on multiple test sets as well as a hold-out external validation set[15]. Of the papers investigated, none contained the full set of verification strategies.

**Table 1.** Results of previous cyclodextrin QSARs.

| QSAR | R2 | Descriptors | Feature selection | Validation |  |  |  |
| --- | --- | --- | --- | --- | --- | --- | --- |
|  |  |  |  | Q2 | *y-rand* | AD | EV |
| Cubist [9] | 0.945 | Pfizer* | ** | ** | ** | Yes | Yes |
| Random forest [9] | 0.912 | Pfizer* | ** | ** | ** | Yes | Yes |
| Partial least squares (PLS) [10] | 0.68 | ChemOffice, SYBYL, Pentacle* | Genetic algorithm | 0.64 | ** | Yes | ** |
| PLS [11] | 0.74 | SYBYL, Pentacle* | Fractional factorial design | 0.75 | Yes | Yes | ** |
| Multiple linear regression (MLR) [12] | 0.943 | ISIS/Draw, CODESSA* | Forward selection | 0.848 |  |  | ** |
| MLR [13] | 0.841 | ISIS/Draw, MOPAC, Web-DRAGON* | Genetic algorithm | 0.821 | Yes | Yes | ** |
| MLR [14] | 0.833 | HyperChem, DRAGON* | Forward selection | 0.826 | Yes | Yes | ** |
| MLR [15] | 0.78 | SYBYL, MOE, AutoDock Tools, BINANA* | Genetic algorithm | 0.82 |  | Yes | ** |
| Artificial neural network [14] | 0.957 | HyperChem, DRAGON* | Forward selection | 0.955 | Yes | Yes | ** |
| Support vector machine [16] | 0.971 | HyperChem, MOE* | Particle swarm | ** | ** | ** | ** |

\*Presence of a paywall, usually due to specialized software that requires a license
\*\*Insufficient verification
None of the models investigated were both fully validated and openly accessible.

In this study, the accuracy and usability of docking and QSARs were compared in order to establish the appropriate framework for the computational design of cyclodextrin based affinity delivery devices. Autodock VINA, an open-source docking program developed by Trott, was used to investigate docking methods[16]. A variety of statistical methods presented in previous cyclodextrin QSARs were also investigated. The performance of QSARs was evaluated on both a standard test set as well as an external validation set to confirm accuracy. Properly evaluating the use of docking and QSARs should improve selection of possible guests for cyclodextrin, reducing the rejection of good candidates (Type II error) and limiting experimental investigation of bad candidates (Type I error).

Finally, though the coded models could be made freely available, understanding the raw script remained a significant obstacle for new users. Additionally, users would have to download multiple files and programs to their own computers, creating potential issues with device compatibility, storage restrictions, processor limitations, etc. To overcome these obstacles and improve accessibility, the models were then integrated into a web application built with the R library “shiny” and then uploaded online. To demonstrate the ease of extendibility of the app and its value in planning drug delivery strategies the results from the QSAR studies were then integrated into a mechanistic model of drug release.

## 2. Materials and Methods

In order to be accessible, the models use only open-source software. Importing experimental data, cleaning data, and creating QSAR models were performed using R in RStudio. Both the coding language and the IDE are freely downloadable and easily accessible on Windows, Mac OS, and Linux. Descriptors were generated with PaDEL, also freely downloadable and open-source. Only the original observations of cyclodextrin complexation energies remain inaccessible to the public, but this does not have any effect on using the models for new predictions.

### I. Dataset

Many of the models in Table 1 work from the same data source, a compilation of α- and β-CD affinities published by Suzuki in 2001[17] (additionally, the sources that cite a different paper by Katritzky ultimately use this same data, as the Katritzky paper cites Suzuki for observations[18]). In addition to Suzuki, we also compiled complexes of α- and β-CD Rekharsky and Inoue and Suzuki[19]. Complexes of γ-CD, missing from the Suzuki dataset and sparse in the Rekharsky and Inoue data, were collected from Connors[20]. Once compiled, the data were cleaned for reliable information, one-to-one cyclodextrin complexes, a temperature of 298 ± 2 K, and a solvent of water with pH 7. To obtain structure-data files (SDFs) of the ligands, the names of the guest molecules were passed through the Chemical Identifier Resolver, a web interface provided by the National Cancer Institute’s Computer-Aided Drug Design Group (NCI/CADD). To handle the data, the R packages tidyverse, data.table, XML, RCurl, and Matrix were used[21–25].

Dataset splitting was performed using the R package caret[26]. First, the cleaned data was split between α-, β-, and γ-CD. Structural and activity outliers in each category were removed. Structural outliers were detected using a statistical method relying on standard deviations of molecular descriptors[27]. For activity outliers, molecules with reported ΔG values greater than 2.5 standard deviations from the mean were removed. Though traditional practice advises classifies outliers as values more than only two standard deviations away, in this case, retaining data points remained a priority and a larger margin was allowed. There were 9, 21, and 11 α-, β-, and γ-CD outliers, respectively. After removal, around 200, 250, and 100 α-, β-, and γ-CD observations remained.

The data was then split into training, testing data and external validation. For each separate cyclodextrin, an external validation set was created from a random 15% subset of the data. To create multiple training and test sets, the remaining modeling data was split with representative resampling of ΔG values into ten different 75:25 train to test data partitions. Though not as advanced as maximum dissimilarity algorithms, this method proved more practical due to the large number of descriptors (over 1,000) generated for each guest molecule. Furthermore, maximum dissimilarity algorithms, when implemented in this instance, had the unfortunate tendency to select highly similar training and test sets, defeating the purpose of creating multiple sets in the first place.

### II. Docking calculations

The process of docking is based on two processes: sampling and scoring[5]. Sampling refers to the capacity to search an active site on a protein, macromolecule or, in this case, affinity host. This can be performed with distance matrices, matching algorithms or incremental construction, multiple copy simultaneous searching, stochastic methods, or any combination of the aforementioned strategies. Scoring calculates the final binding affinity between the guest and host and can be dependent on force-field, empirical, or knowledge-based calculations. Docking generally involves the use of a host and a guest molecule which can be either rigid or flexible. Three types of conformation exist: rigid-rigid, rigid-flexible and flexible-flexible. In this paper we use AutoDock Vina, a version of AutoDock that uses Monte Carlo stochastic sampling coupled with a force field based scoring function from a resample of a drug like database to derive its weighted parameters. Vina in particular uses a flexible drug guest and a rigid cyclodextrin host, although it allows side chain mobility when docking ligands onto proteins.

The PyRx Virtual Screening Tool provides a variety of services, including molecular energy minimization, docking calculation, and visualization of molecules. PyRx version 0.8 was used here [28]. To begin, all guest molecules went through energy minimization to determine the most likely atomic configurations. AutoDock Vina, integrated within PyRx, calculated the change in Gibbs free energy (kcal/mol). We tested the effect on the docking process of changes in the search space, search exhaustiveness, and scoring force field type.

### III. Descriptor Generation

The open source software PaDEL-Descriptor calculated over 1000 descriptors for the remaining molecules, including fingerprints, structural details, and physical properties[29]. Additionally, PaDEL-Descriptor removed salts and minimized the energy of inputted files using an MM2 force field. To improve model interpretability, more abstract predictors, such as those related to eigenvalues for molecular matrices or autocorrelation, were excluded from calculation. The elimination of these descriptors did not produce any noticeable effect on final model accuracy and made feature selection less resource intensive.

### IV. Feature selection

Recursive feature elimination (RFE), implemented with caret, was used to subset the predictors used for model-building (Kuhn 2018). Using this method, a random forest model is created using all available descriptors. Once trained, the relative importances of the predictors are calculated and differently sized subsets (defined by the user) of variables are selected to create and evaluate new models. The best combination of predictors is then returned by the model. RFE was performed on each of the ten train-test splits. The predictors determined to be useful for all folds were saved and used for tuning and training the models. This resulted in 13 variables for α-CD, 16 variables for β-CD, and 39 variables for γ-CD.

### V. QSAR development

We investigated the accuracy of several models that appeared in previous attempts at cyclodextrin QSARS (Table 1), including Cubist models, generalized linear models (GLM or GLMNet), random forests, partial least squares models, and support vector machines. Additionally, two QSAR methods not previously published for cyclodextrin – multivariate adaptive regression splines (MARS) and gradient-boosted models – were created and evaluated. Model building was accomplished with R-packages Cubist, glmnet, randomForest, pls, e1071, earth, and gbm, respectively [30–34].

Cross-validation was used to determine ideal tuning parameters for each QSAR. For faster QSARs – such as generalized linear models (GLM) and partial least squares (PLS) – tuning was performed using 10-fold cross validation. For more resource-intensive models or models with large parameter spaces – such as random forests, Cubist and support vector machines (SVM)– only 5 folds were used. Optimized models, QSARs built with the tuned parameters and trained on the entire training set, were used to predict the test for each combination of test and training set. Further fine tuning was also performed at this step. The model that produced the lowest root-mean square error (RMSE) and highest R^2^ (or an otherwise most ideal combination) on all the test sets became the final model, i.e., the model saved for future use. Furthermore, the models were evaluated according to Tropsha and Golbraikh standards for QSARs[35]. Although R^2^ and RMSE can be useful for generalizing predictive capacity, they may be misleading in certain cases, necessitating stricter additional standards of evaluation. As an additional test of reproducibility, the final models were used in ensemble to predict the values of the external validation set. Because this dataset was withheld from the entire model training process, the external validation set served to simulate model performance on new data.

### VI. Applicability domain

Applicability domain describes the range of molecules where the model can be expected to generate reliable predictions. A new molecule outside of the applicability domain is structurally quite different from the set of data the model was trained on, and thus a prediction will rely on extrapolation and may not be accurate. The applicability domain of the models was determined with the same method used to detect outliers when cleaning the dataset[27].

### VII. Y-randomization

Y-randomization was used to further verify the significance of the results. Many advanced QSAR methods are powerful enough to model data off of noise, so y-randomization ensures that the modelling process produces results significantly more accurate than what could be obtained by chance. Randomization can be achieved by permutation (randomly changing the positions of observed values) or random number generation (replacing observed values with completely new data). Different combinations of permutation and/or random generation yields five different modes of y-randomization to investigate: 1) original ΔG values vs. randomly generated descriptors, 2) permuted ΔG vs. original descriptors, 3) random ΔG vs. original descriptors, 4) random ΔG vs. random descriptors, and 5) permuted ΔG vs. random descriptors. (Combinations including permutation of descriptors are not included because the large number of predictors in QSARs renders the effects of such a process virtually indistinguishable from random number generation.) However, because the y-randomization process is extremely resource-intensive (as each mode requires that several randomized iterations undergo the modeling process), only mode 1, the most common interpretation of y-randomization, was investigated[36]. The observed ΔG values were randomly assigned to guest molecules, and the entire model refitting process was re-done, from feature selection to external validation.

### VIII. Creating an App

Using R’s “shiny” package, most of the process of running the QSAR could be implemented in a web app. The app was split into three main pages: Download, Upload, and Explore. “Download” accesses Chemical Identifier Resolver and obtain SDFs. The page also draws the obtained molecule using the package “ChemmineR”, allowing the user to check that the SDF is accurate. “Upload” implements the QSARs after the user provides the app with a CSV of the descriptors from PaDEL-descriptor. After calculating the affinity and analyzing the applicability domain of the molecules, the user is provided with both a graph and a table of the results. The third page “Explore”, stores the results of using the ensemble on FDA-approved drugs, as obtained from the annual publication “Orange Book: Approved Drug Products with Therapeutic Equivalence Evaluations.” [40]

### IX. Modeling release curves

Partial differential equations that model drug diffusion were solved using the R package deSolve[37] using the method of lines as previously done by Fu[2]. In the model, the release media was assumed to be water and the delivery system was assumed to be flat, thin circular cyclodextrin disc. The boundary condition between the polymer and the media was approached as described by Linge [41]. Diffusivities of drug molecules were calculated from molecular weight and viscosity using a modified Stokes-Einstein-Sutherland equation, as done by Vulic [38].

## 4. Results and discussion

### I. Performance of docking

When predicting on the entire cleaned dataset (all modeling data, which includes the training, testing, and external validation set), AutoDock Vina yielded an R^2^ of 0.18 and a RMSE of 5.00 kJ/mol (Fig 2). Of the 547 cleaned complexes, docking provided calculations for 458, failing to provide data on 89 complexes. Adjusting settings in Vina, such as the minimization algorithm size of the steps in the calculation, did not yield significant differences in accuracy. In comparison, the affinity of only around 40 cleaned molecules could not be obtained by the ensemble QSAR. In these cases, the withheld molecules were determined to be outliers, and the actual QSAR model could still be used to predict a value.

**Fig 2.**
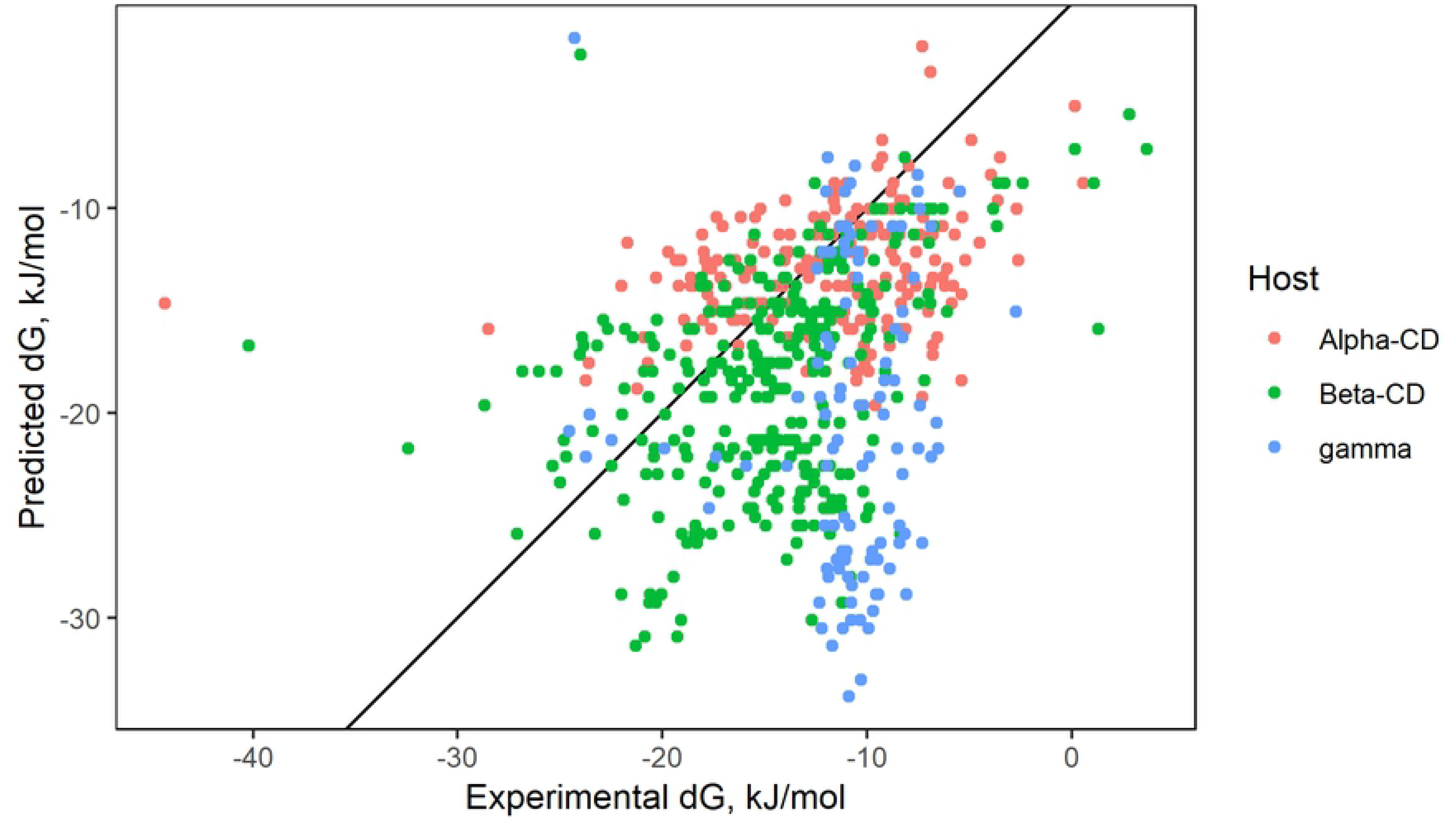
Results of PyRx docking.

### II. Performance of QSARs

The results of predicting on the test data for each QSAR and cyclodextrin type are markedly higher than docking (with the exception of γ-CD), reaching an R^2^ of around 0.5 to 0.7, as seen in Table 2 and Fig 3. The reported R^2^ for each QSAR type is calculated from an average of the performance of the model on all test splits. Additionally, Table 2 contains information on verification of all the QSAR types (3-VII Validation methods). Of the types investigated, only PLS and GLMNet failed to pass the salvo of verification criteria, both falling short of attaining an R^2^ of 0.6.

**Table 2.**
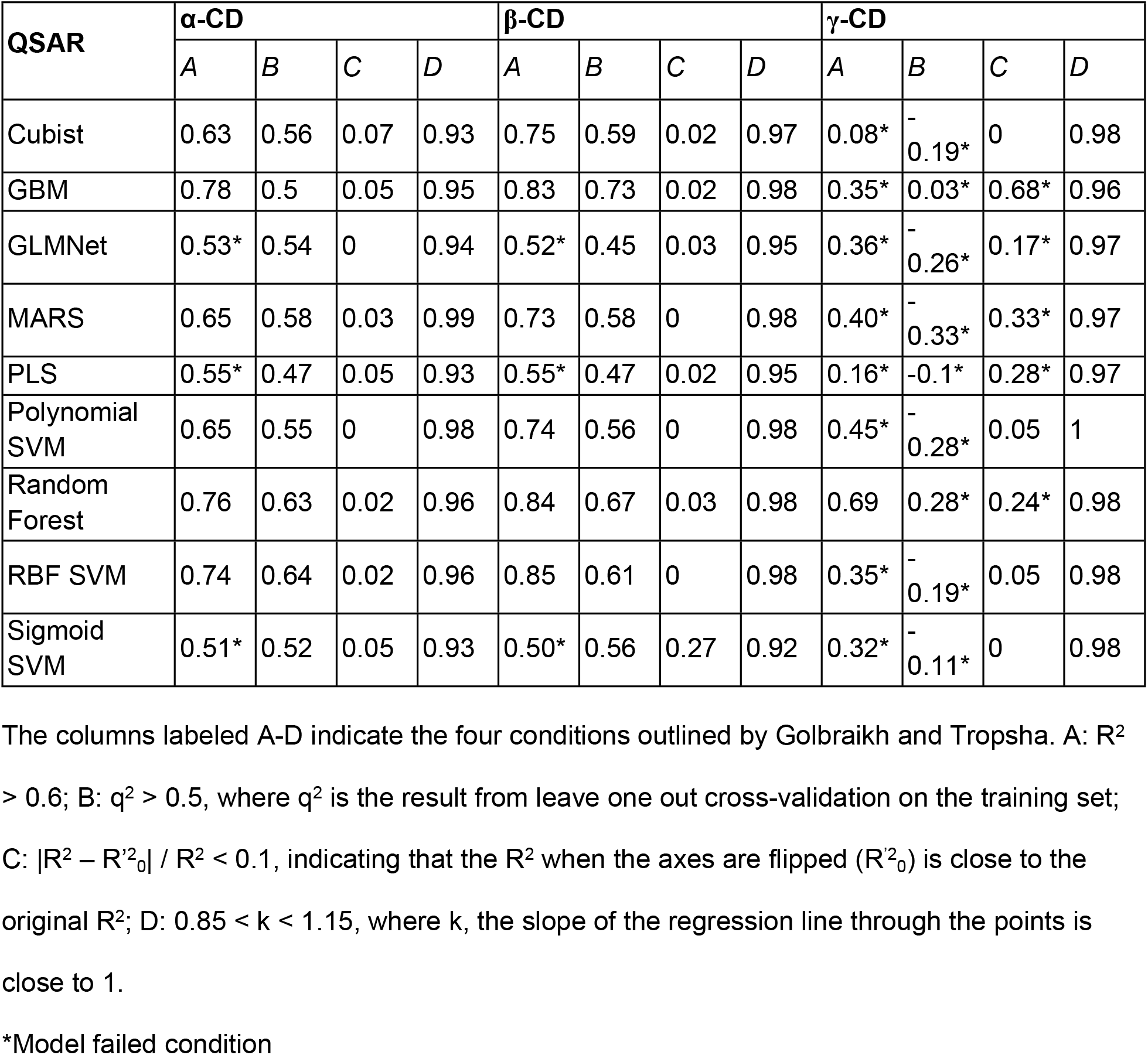
Evaluation of QSARs on test sets.

**Fig 3.**
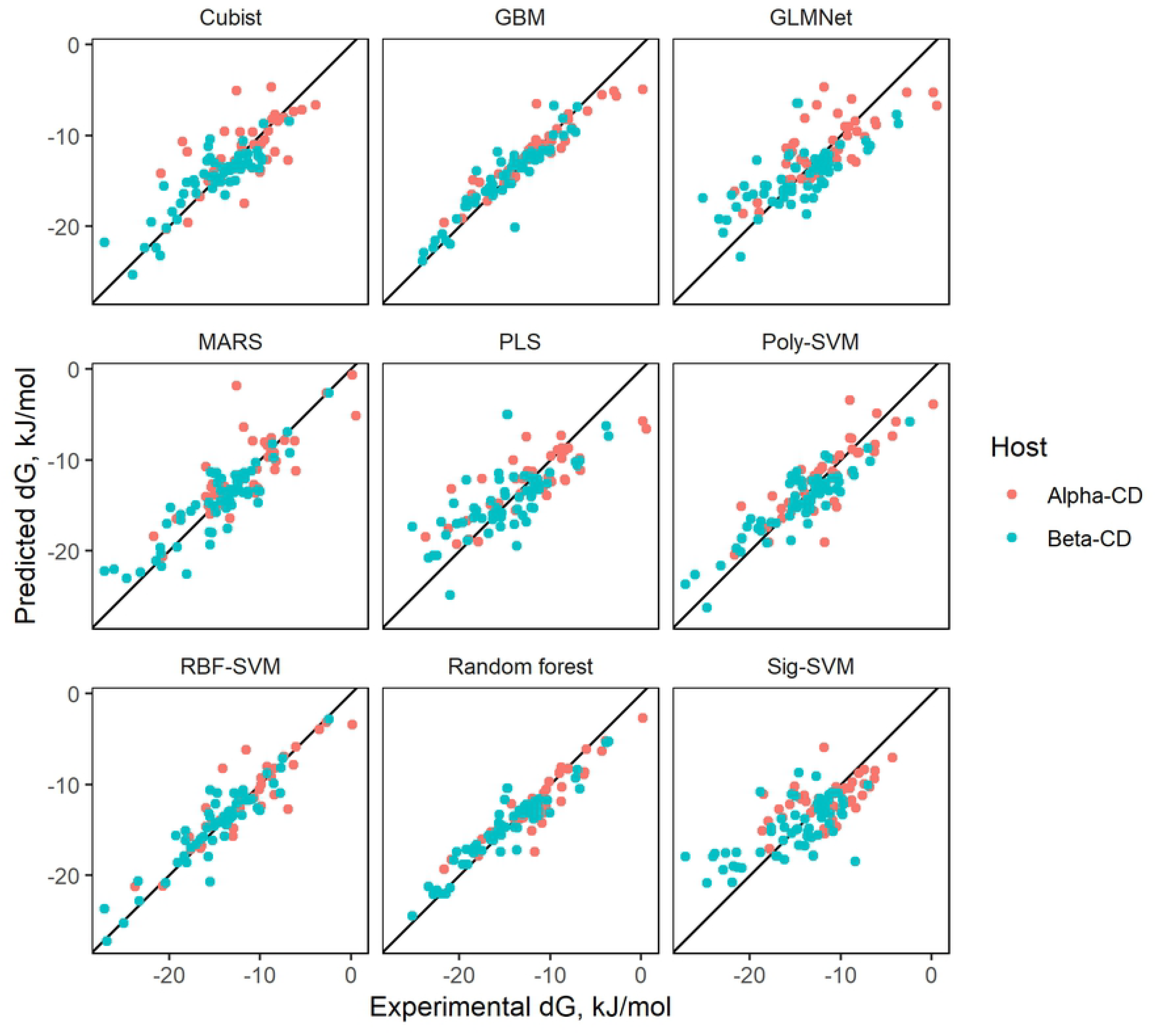
Results of QSARs on test sets.

In terms of reliability, most models were able to handle the available data well, providing calculations for all provided molecules. Only the Random Forest and Cubist models failed to calculate the affinity of some molecules, possibly due to being based around decision-trees. The algorithm underlying both models attempts to draw predictions by categorizing entries based on their features. If they encounter a molecules entirely different from the data they trained on, the models may fail to create a prediction. Advantageously for our approach, the failure to calculate some values becomes less important where models are combined in an ensemble where the final prediction is averaged over many models.

The results of ensemble prediction (averaging the results of many different QSARs) can be seen in Figure 4. While α- and β-CD models managed to reach moderately high predictive performance metrics, unfortunately, all γ-CD models lacked usable predictive power.

**Fig 4.**
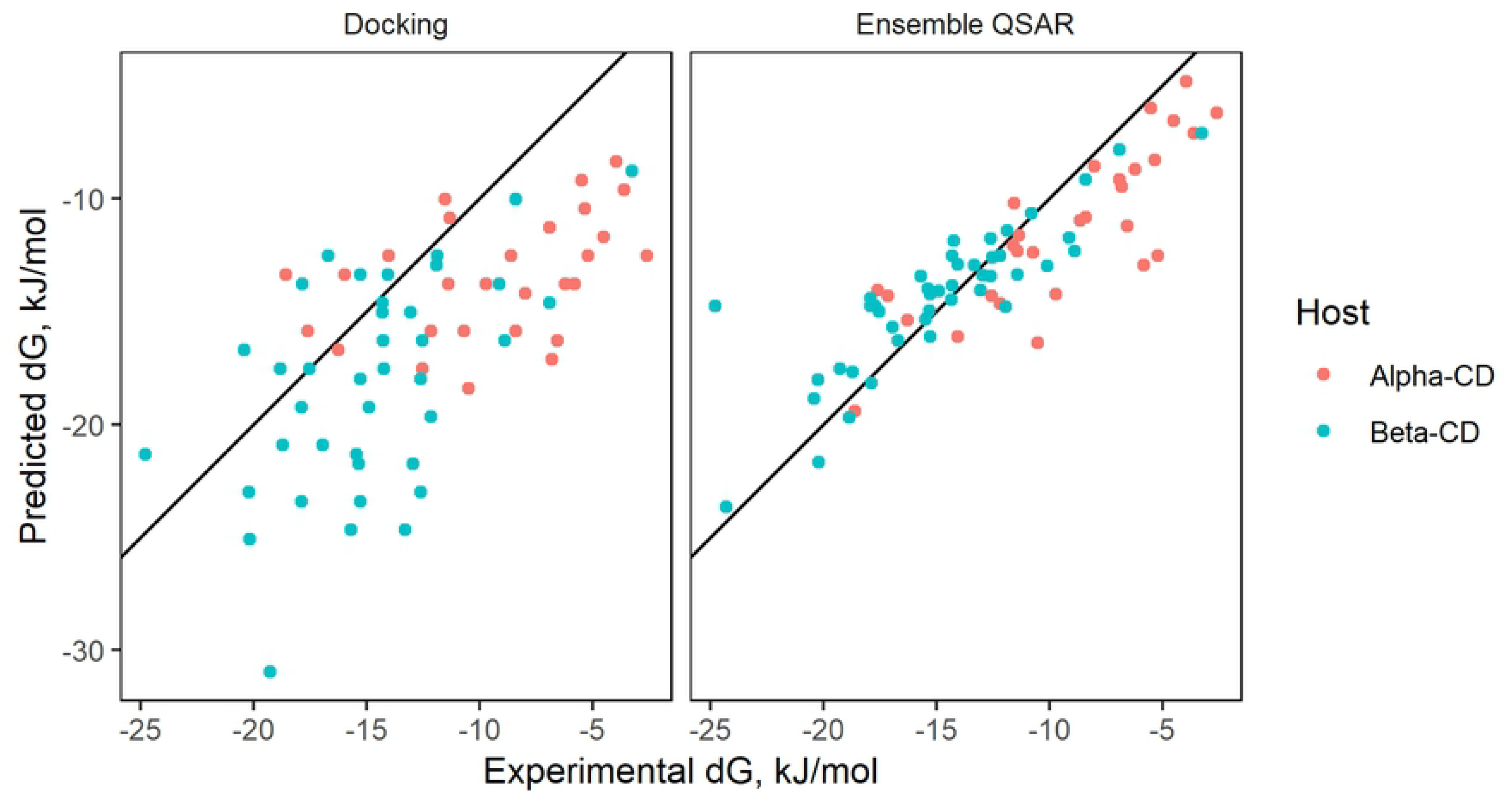
QSAR ensemble prediction.

The models passing the verification in Table 2 were further verified using y-randomization. To ensure accuracy was not the result of the models building off of noise, 25 different permutations of ΔG values were created. All Q^2^ values of the models created from the original data were calculated to lie well outside 3 standard deviations of the mean Q^2^ of the randomized data. Additionally, the R^2^ values of the ensemble QSARs were significantly greater than the R^2^ values obtained from the ensemble models created from permuted data (means of 0.021 and 0.027 and standard deviations of 0.011 and 0.016 for α- and β-CD, respectively).

### III. Variable importance

Interpretability of a model provides a rough check if a model is calculating off of random noise or if the model is drawing logical calculations from physical properties to molecular behavior. Each model, due to differences in statistical algorithms and approaches, has differing levels of interpretability. GLM, being similar to linear models, have easily accessible coefficients associated with each predictor, so the relative impact of each factor can be compared with reasonable confidence. Cubist models, on the other hand, tend to be difficult to interpret as variables are processed through multiple levels of decision trees.

Evaluation for the relative importance of variables are shown in Figure 5. Random forest was the only QSAR type with a pre-packaged importance function for variable analysis. PLS variables were analyzed using a function obtainable from Bjørn-Helge Mevik[39]. Max Kuhn’s caret package was used to evaluate GLMNet, the two SVM kernels, and Cubist. Unfortunately, caret was unable to process the final models for GLMNet and SVM, and the reported variable importance values were actually derived from models created within caret’s “train” function, and are thus slightly different from the models saved in the ensemble. To determine importance, the “train” function removes a variable, rebuilds the model, and analyzes the effect on accuracy. The more important a variable, the larger the drop in accuracy. After each variable has been tested, the function can then rank the importance of the descriptors.

**Fig 5.**
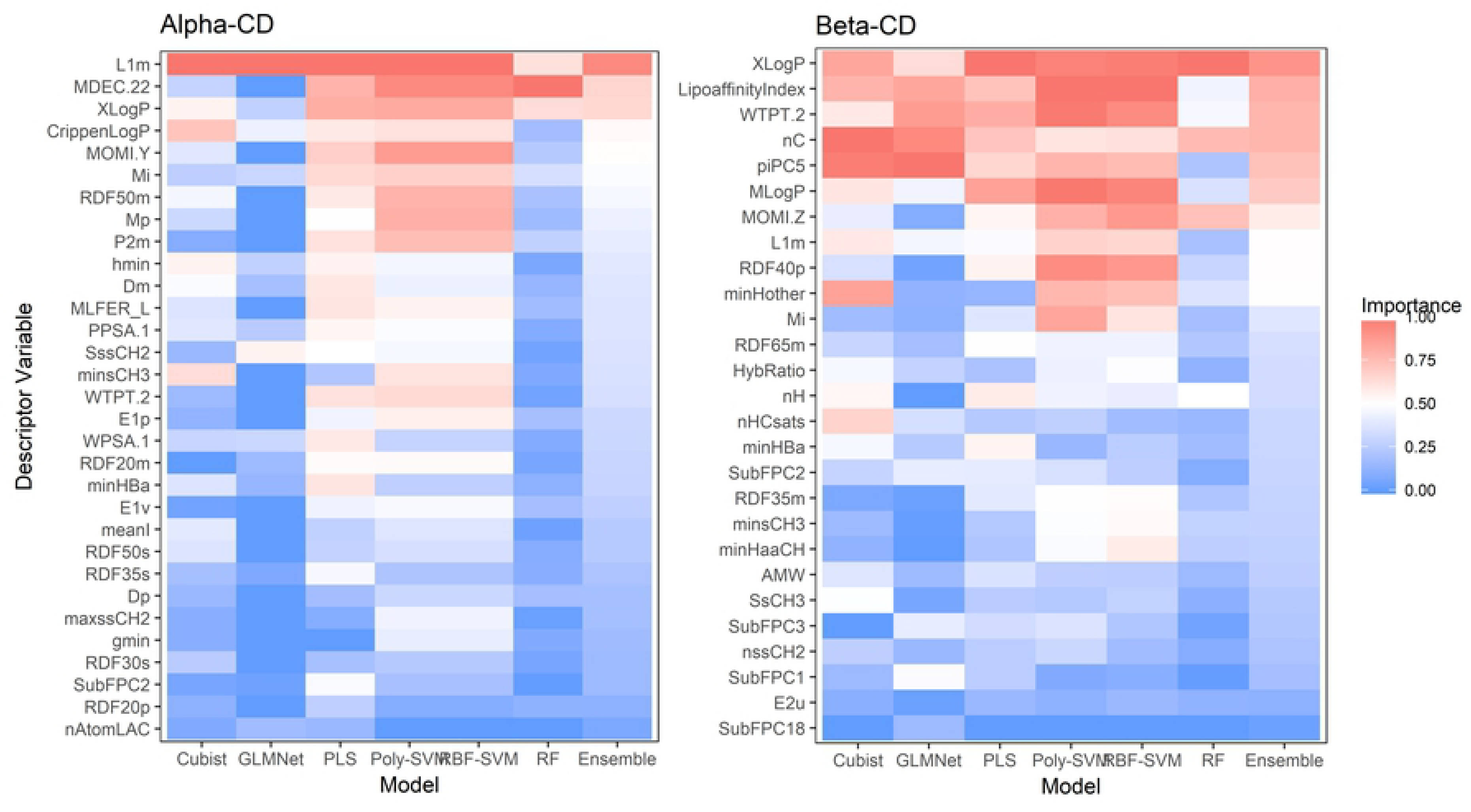
Variable importance.

**Fig 6.**
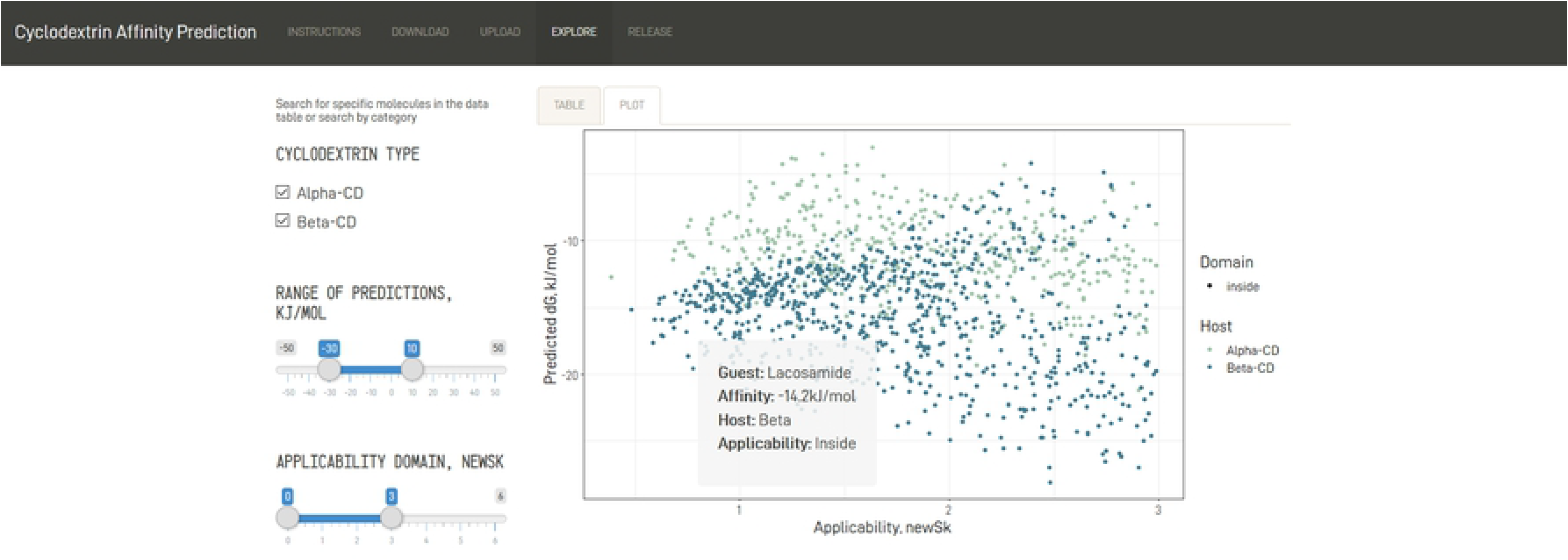
User interface of the Shiny App.

**Fig 7.**
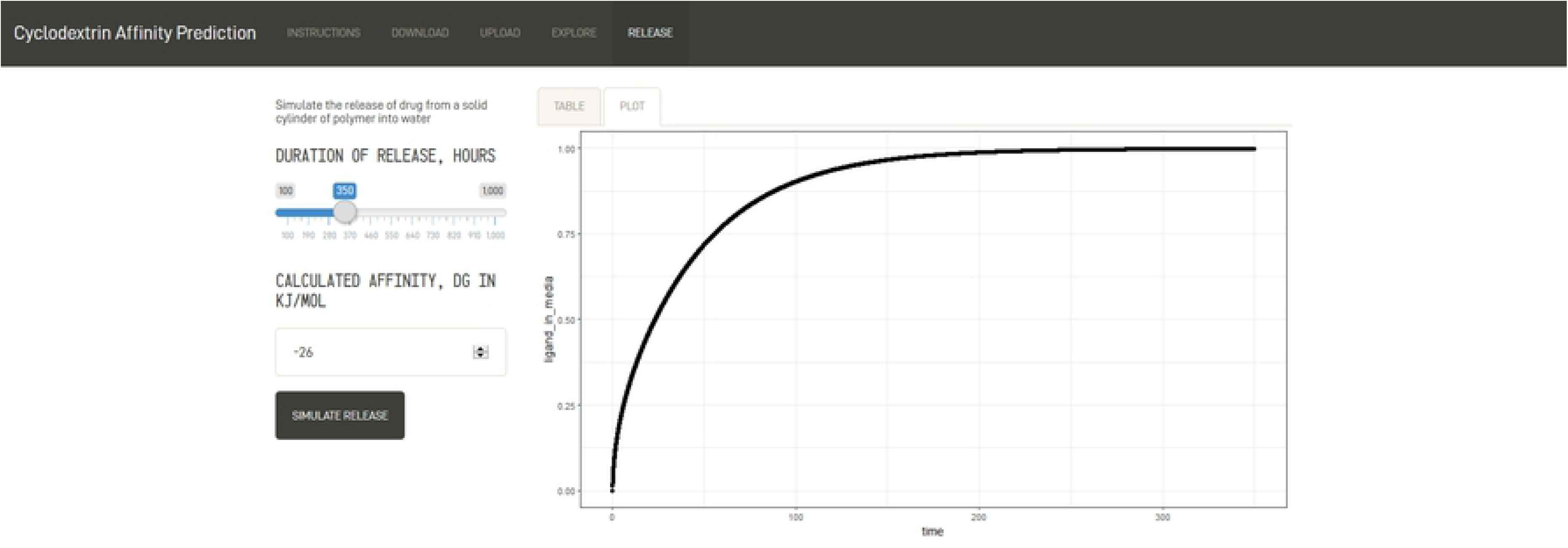
Drug release curves.

For β-CD, XLogP, a measure of lipophilicity, appears to be important for all models, consistent with how the structure of cyclodextrin allows for easier complexation with small hydrophobic drugs. The same reasoning can be extended to LipoAffinityIndex and MLogP, additional approaches to quantifying lipophilicity. The number of carbons, nC, is also consistently important, possibly due to a relationship with molecule size. WTPT-2 is the PaDEL weighted path descriptor divided by the number of atoms, and also may be important due to encoding information on molecular size.

However, not all the variables can be linked to set chemical properties. SpMax and SpMin relates to eigenvalues of a modified connectivity matrix, a numerical representation of atomic and molecular bonds, and may not be associated with any interpretable physical property (the same analysis can also be used for GATS predictors). To aid interpretability, building a model with predictors easily attributed to physical or chemical properties may be advised. The extent to which interpretability should trade off with accuracy remains in question. Our findings go in accordance to those in the general literature from table 1 were lipophilicity tends to highly influence model output.

### IV. Web Application and FDA Database

After collecting a list of FDA-approved drugs and drug combinations from the Orange Book, an annual publication listing all approved pharmaceuticals [cite], the names were cleaned for individual active compounds. In total, 1401 unique molecules could be extracted. Of these, 1116 could be downloaded from Cactus and 1031 could be processed by PaDEL. Many of the molecules that could not be analyzed by PaDEL would have proven impractical for cyclodextrin delivery, such as simple ionic salts (e.g., potassium chloride), or large molecules made of more than 100 atoms. Running the remaining guests through applicability domain analysis yielded 638 molecules, 45.5% of the original set. While less than half of FDA-approved drugs could pass through the model, the 600 available guests spans a wide range of properties and uses, allowing the page to be useful for candidate selection.

Though the app could be uploaded online through shinyapps.io, server time limitations on the account hosting the app make it impractical for usage by a large number of individuals simultaneously. In order to run the app for more than a few hours, such as with screening a large dataset of molecules, the code would have to be downloaded through GitHub. In addition, the user would need to download the R libraries and the IDE RStudio, potentially negating the goal of creating an accessible, intuitive interface. The “Explore” page partially alleviates this obstacle, as it allows the user to perform a quick search of a pre-predicted affinity rather than spend time downloading the structure file, launching PaDEL, and running the QSAR.

### V. Drug release module

To demonstrate the extensibility of the shiny app and its value in the design of drug delivery strategies the results of the QSAR predictions can be fed into a mechanistic model of drug delivery. The results demonstrate the ranges of values expected from the strongest affinity binding predictions and from the weakest. As expected, strong predictions produce much slower release profiles and weak predictions produce faster release profiles. Conservation of mass was verified as the sum of all mass within the system from the polymer and media compartment across all times as a test of the implementation. Notably, the implementation in R required a modification of the method of lines for appropriate modeling of the polymer to liquid media interface.[41] Future efforts in creating new modules could explore substructure searching to identify alternative strategies for weak binders or drugs that demonstrate unsuitable release profiles. It is expected that not all drugs will follow this simplistic model of drug release. For those cases our lab has built a whole suite of approaches to alter elution rates such as a wide range of formulations, supramolecular interactions, Schiff-base formation and multi-arm PEG substitutions.

## 5. Conclusions

In predicting the binding affinity of cyclodextrin with small drug molecules, QSARs such as Cubist, GBM, MARS, random forest, and SVM models can be created using accessible open-source software. These models outperform available molecular docking software in both accuracy and time consumption and pass statistical verification of reliability. The additional accuracy afforded by QSARs can be integrated into the previously published mechanistic model for predicting drug release curves for candidate molecules. This would both help narrow down candidates for affinity-based drug delivery using cyclodextrins, as well as help advise drug loading configurations to tailor release rate from a delivery system for a given biomedical application. Additionally, similar strategies could model other molecular interactions and allow application of computational approaches for higher-throughput research strategies in drug delivery, an inherently empirical field. Approaches like the one presented here will move drug delivery closer to a truly integrated approach in drug discovery.

## Acknowledgements

The authors would like to acknowledge the Hathaway Brown School and Dr. Crystal Miller for enabling Alison Xin’s contributions through the Science Research & Engineering Program.

